# Larvicidal Potential of *Cola nitida* Endophytic Bacteria Against Insecticide-Resistant *Anopheles gambiae*

**DOI:** 10.64898/2026.08.03.742416

**Authors:** Jean Baptiste Hzounda Fokou, Steve Henri Voundi Olugu, Cyrille Ndo, Alex Kevin Tako Djimefo, Julienne Victoire Azienwi, Angelique Nghariegam Nsangou, Jean Emmanuel Mbosso Teinkela, François Eya’ane Meva

## Abstract

**Background:** Innovative vector control strategies are urgently needed to combat malaria transmission by *Anopheles gambiae*, given the limitations of current insecticides—namely resistance, human toxicity, environmental damage, and high costs. This study evaluated the larvicidal efficacy of endophytic bacteria isolated from *Cola nitida* against third-instar larvae of *Anopheles gambiae*.

**Methods:** Endophytic bacteria were isolated from nine *Cola nitida* plant parts (leaves, roots, stems, fruits, flowers, stem bark, branches, root bark, whole plant extracts) using standard surface sterilization and culturing protocols. Thirty-two morphologically distinct strains were purified and screened for larvicidal activity against third-instar *An. gambiae* larvae using the WHO-recommended turbidity method at McFarland 4 standard. Six active isolates underwent dose-response testing (McFarland 0.5-4) with mortality recorded at 24 and 48 hours. Environmental safety was assessed via *Lemna minor* growth inhibition, and molecular identification employed the API 20E biochemical gallery.

**Results:** Screening identified six strains with significant larvicidal activity (LC₅₀ < McFarland 2). At McFarland 2 concentration, all six strains achieved >50% mortality within 24-48 hours, with *Lemna minor* EC₅₀ >100% indicating minimal phytotoxicity. API 20E identification (96-99.99% similarity) revealed three strains (6164, 6211, 6501) matching *Photobacterium damselae*, two (6186-1, 6512-1) matching *Salmonella enterica* subsp. *arizonae*, and one (6600) matching *Pseudomonas* sp., confirmed by distinctive biochemical profiles.

**Conclusion:** This study provides the first evidence of potent, environmentally safe larvicidal activity from *Cola nitida* endophytic bacteria against *Anopheles gambiae*. These strains represent promising biocontrol candidates to address insecticide resistance, offering a sustainable alternative for malaria vector management in endemic regions.

## Introduction

Vector-borne diseases are human illnesses caused by parasites, viruses, or bacteria and transmitted by vectors such as mosquitoes, ticks, and flies. These diseases account for over 17% of all infectious diseases, leading to more than 700,000 deaths annually, with the highest burden in tropical and subtropical regions disproportionately affecting the world’s poorest populations[1]. Among these, malaria, transmitted by female *Anopheles* mosquitoes, poses a major public health threat. In 2023, approximately 263 million malaria cases were reported globally across 85 countries, resulting in approximately 597,000 deaths, with Sub-Saharan Africa accounting for 94% of cases and 95% of fatalities.[2–5] In Cameroon, the situation is particularly alarming, with 3 million confirmed cases and 1,756 malaria-related deaths in 2023, accounting for 11% of all health facility deaths; 78% among children under five years old[6].

Eliminating malaria remains one of the most ambitious goals in global public health, and achieving it will depend, in large part, on our ability to control the mosquitoes that spread it [1, 7]. At its core, vector control aims to reduce or suppress *Anopheles* mosquito populations enough to interrupt transmission from one person to another.

The tools most widely used today fall into two broad categories: chemical interventions including insecticides and larvicides and physical or environmental measures, including the drainage of stagnant water bodies and the clearance of dense vegetation where mosquitoes breed and rest[1, 8–10]. These methods have formed the backbone of malaria control programmes for decades and, in many settings, continue to deliver meaningful results.

Yet the limitations of these approaches are becoming increasingly difficult to ignore. Mosquito populations in many regions have developed resistance to commonly used insecticides, while concerns over toxicity to human health and collateral damage to ecosystems have raised serious questions about their long-term sustainability. Add to this the considerable financial cost of large-scale implementation, and it becomes clear that conventional methods alone are unlikely to be enough[1, 9–13]. There is, therefore, a persistent need for new and smarter approaches capable of keeping mosquito populations in check while reducing the risks of resistance, ecological harm, and toxicity to non-target species.

Biological control strategies against mosquitoes draw on naturally occurring agents including pathogens and organisms that live in close association with mosquitoes. Several real-world applications have demonstrated the value of these approaches. Community-based programmes introducing cyclopoid copepods (*Mesocyclops longisetus*) and various species of fish into mosquito breeding sites have successfully reduced populations of dengue vectors [14, 15]. That said, the long-term effectiveness of these interventions depends heavily on the characteristics of the breeding sites themselves and how productive those environments remain over time.

Beyond these approaches, recent years have seen growing interest in technologies that rely on “modified mosquitoes” to suppress wild populations. These include genetic modifications as the introduction of dominant lethal genes (RIDL) or RNA interference (RNAi) immunity genes. as well as non-genetic methods like the sterile insect technique (SIT), which uses radiation to render males infertile, and the incompatible insect technique (IIT), which exploits naturally occurring bacterial incompatibility. Importantly, SIT and IIT do not involve genetic modification and are therefore not classified as genetically modified organism (GMO) technologies[16].

On the biorational side, formulations based on *Bacillus thuringiensis* var. *israelensis* (Bti) have proven highly effective as bio-larvicides against both malaria and dengue mosquito vectors [17, 18]. These products target larvae selectively, making them a practical and environmentally friendly tool in integrated vector management programmes.

Endophytes, microorganisms that colonize plant tissues without causing disease, have emerged as promising tools in the biological control of human vector-borne diseases [19–21]. Both fungal and bacterial endophytes produce a range of bioactive metabolites that can act as larvicidal and antifeedant agents against insect vectors such as mosquitoes, thereby reducing the survival and development of larvae that transmit pathogens like malaria, dengue, and chikungunya viruses [19–21]. Endophytic fungi isolated from medicinal plants, including *Ocimum sanctum*, *Vitex negundo*, and *Azadirachta indica*, have shown significant larvicidal activity against *Anopheles* and *Aedes* mosquito larvae, often at low concentrations and with toxicity comparable to or higher than standard plant extracts[20, 22, 23]. Bacterial endophytes, particularly strains of *Bacillus* and *Pseudomonas*, further enhance this potential by synthesizing insecticidal proteins and secondary compounds that disrupt larval physiology and feeding behaviour [24–26].

To the best of our reading, there is no report on larvicidal effect of endophytes from Cola nitida. This study aimed at evaluating the effect of endophyte on third Larval stage in a coculture model.

## Material and Methods

### Study type

This was an experimental study.

### Study location

This study was carried out at the Laboratory for Pharmacology of the Faculty of Medicine and Pharmaceutical Sciences at the University of Douala.

### Period of study

This study ran for a period of 8 months from November 2024 to June 2025.

### Study authorisation and approval

The study was granted the authorisation number 90/05 AR/25/UD/FMSP/CDAASSR/mac by the head of the academic affairs division of the faculty of Medicine and Pharmaceutical Sciences, the University of Douala.

The ethical clearance number 4904/IEC-Udo/04/2025/M was obtained from the institutional ethical clearance committee of the University of Douala.

## Study materials

### Plant material

The parts of the plant *Cola nitida* used for this study include the roots, root bark, trunk, branches, and leaves. were harvested in the Littoral region of Cameroon at Njombe Penja and identified at Yaoundé National Herbarium under identification number 48651 SRF Cam.

### Organisms used

#### Obtaining and rearing of Anopheles gambiae larvae

To assess the larvicidal activity and lethality testing, *Anopheles gambiae* eggs were graciously donated by the Centre for Research in Infectious Diseases (CRID), from Professor NDO Cyrille.

These eggs were reared according to the WHO guidelines for laboratory and field testing of mosquito larvicides.

As eggs hatched, the larvae were transferred to shallow trays containing mineral water. The aim was to create a population of same instar larvae per container. The containers were kept at room temperature. The amount of food was kept low to avoid strong bacterial growth (which kills the larvae), increasing food provision as the larvae grow. Several feeds at intervals of one or two days and daily observation of the larvae. A homogeneous population of third instar were obtained five to seven days later.

#### Obtention and culture of Lemna minor

Healthy and fresh Lemna minor plants (with green fronds) used for this experiment were collected in the Yassa neighbourhood in Douala. These plants were acclimatized in a modified Lemna growth medium under natural laboratory bench conditions.

## Methods

### Preparation of crude extracts from the endophytes

The methods used here follow the Pharmacology Laboratory protocol as described by Hzounda *et al.[27]*

#### Identification and harvest of the plant

The plant *Cola nitida* was harvested in the Littoral region, specifically from the Njombe-Penja community in the Mungo division and immediately transported to the Pharmacology Laboratory of FMSP-UDo. It was authenticated at the Cameroon National Herbarium by comparing it with a laboratory sample registered under reference number 48651 SRF Cam.

#### Post-harvest surface sterilization of the plant

The surface sterilization method proposed earlier in the literature [27–29].After the harvest, the plant was brought to the laboratory, the various plant tissues were separated (root bark and roots, stem back and stem, branch bark and branch) and washed thoroughly in running water to remove soil and epiphytes. The plant was sorted by checking the parts of the plant: the roots, trunk, branches, leaves were checked to eliminate parts with any damage or discolorations. The healthy plants left were then separated by cutting into small fragments of about 1 cm x 1 cm using a sterile bistoury blade. The explants obtained were surface sterilized by immersion into sterile ethanol at 70 degrees for 2 minutes. The immersed plant pieces were next rinsed in sterile physiological water 3 times to remove disinfectant. They were further immersed in 1% sodium hypochlorite for 1 minute and then rinsed as before. The sterile samples were dried under aseptic conditions around a Bunsen burner. To validate the effectiveness of the surface sterilization, the last rinsing water was inoculated on NA and incubated at 30 degrees Celsius for 24 hours and observed for any growth.

#### Isolation and purification of endophytic bacteria

The isolation and purification of endophytes was done using plant tissue culture and confirmed by microscopy. The culture media were prepared according to the manufacturer’s recommendations. The surface-sterilized ex-plants pieces were seeded directly onto agar plates by placing single plant tissue fragments evenly on an agar plate. Each of the four parts had its pieces spaciously seeded unto; Mueller Hinton Agar and Nutrient Agar + 0.1mg/ml ketoconazole for promoting endophytic bacterial growth and incubated for 30° C for 48 hours. After incubation, the visible bacteria growing out of the plant tissue were cultured unto new MHA plates coded by plants parts and incubated. They were repeatedly streaked in plate count agar every 24 hours until obtention of pure colonies in each plate, confirmed morphologically by macroscopy and microscopy.

These pure isolates were coded with NPMRU6XXX. Micro tubes containing 500 μL of MHB each were prepared, labeled and sterilized. About 2 loops of fresh endophytic bacteria were introduced into corresponding micro tubes then incubated for 24hrs. After respective incubation, 500 μL of sterilized glycerol was added and the tubes were stored at -20 degree Celsius for long term. Another sample was stored at 4 °C, suitable for short-term handling.

### Larvicidal potency evaluation

#### Preliminary screening

The larvicidal effect of isolated endophytic bacteria from Cola nitida was assessed using the standard procedure established by the World Health Organization[16]. To prepare various test suspensions, we used the McFarland 4 standard as a reference. This involves visually comparing the turbidity of our bacterial suspension, to that of the standard which has a concentration of 12 × 10^8^ CFU/mL. Each test solution was placed in 12-well plates, into which 10 Anopheles larvae were introduced. 0.9% NaCl solution served as the negative control. After 48 hours of exposure, the number of dead larvae was counted, and the percentage of mortality reported.

#### Determination of the optimal dose

After confirming our endophytic bacteria possessed certain larvicidal activity, we further tested at varying Mc Farland concentrations, so as to obtain the exact bacterial concentration responsible for this activity (optimal dose) as well as the toxic dose. Following the same procedure as above but at different Mc Farland concentrations; which were 0.5, 2 and 4, we introduced 5 larvae per test solution. After 24hours the larvae are monitored, and then larval mortality is recorded, also a dose-response relationship is evaluated using the standard procedure established by the World Health Organization[16].

### Environmental toxicity testing

#### Lemna minor growth inhibition test

To evaluate the ecological safety of the bacterial isolates derived from *Cola nitida*, a seven-day toxicity assay was conducted using *Lemna minor* (commonly known as duckweed) in accordance with the OECD Test Guideline 221[30]. This method assesses the impact of chemical and biological agents on the vegetative growth of aquatic plants and it is widely recognized for its sensitivity and reproducibility.

#### Test Organism and experimental set up

Healthy and fresh *Lemna minor* plants (with green fronds) used for this experiment were collected in the Yassa neighbourhood in Douala. These plants were acclimatized in a modified Lemna growth medium under controlled laboratory conditions. The experiment was carried out at a relatively constant ambient temperature of 24 ± 2 °C continuous bright white fluorescent light equivalent to 6500-10000 lux (which is similar to natural daylight near a window).

The experimental setup involved transferring 10 *Lemna minor* plants having 2 fronds each into plastic cups, each containing 5ml of our test solutions. The test groups consisted of concentrations of bacterial extracts respectively 2 and 4 Mcfarland standard. The positive control group containing only the nutrient medium. The test followed a static design.

#### Parameter Observation

Throughout the experiment, plant reponses were monitored on days 0,3 and 7. The primary endpoint was the number of fronds, while secondary endpoints included changes in frond colour (chlorosis).

### Endophytic bacteria identification

#### Morphologic characterization

Endophytes exhibiting the desired antibacterial profile underwent culture and subsequent characterization through a three-step approach. Colonies grown on Mueller-Hinton agar were first examined macroscopically, with attention to color, texture, and uniformity across the plate. Individual isolates were then mounted for light microscopy, where shape and motility revealed key morphological distinctions between strains. Species-level identification followed, relying on the API 20E gallery to resolve biochemical profiles and assign taxonomic identity to each isolate.

#### Endophytic bacteria identification

Endophytes of interest were cultured on MHA and assessed by colony color, appearance, and uniformity, then examined microscopically for cell shape and motility. Species identification used the API 20E system, which comprises 20 miniaturized biochemical assays on a plastic strip for Enterobacteriaceae and other Gram-negative bacteria. A suspension in 0.9% NaCl was standardized to 0.5 McFarland and inoculated into each microtube; mineral oil covered anaerobic tests (ADH, LDC, ODC, H2S, URE). Strips incubated at 36 ± 2 °C for 18–24 h with humidity control. After incubation, reagents (ferric chloride, Kovac’s, VP reagents) were added, colour reactions recorded, and the resulting profile converted to a numeric identification code.

Biochemical test functions

Each test in the API 20E strip is listed with its purpose in the table (Table 1) below:

**Table 1:** Each biochemical test and its function.

| Test Code | Function |
| --- | --- |
| **ONPG** | Tests for β-galactosidase |
| **ADH** | Arginine dihydrolase activity |
| **LDC** | Lysine decarboxylase activity |
| **ODC** | Ornithine decarboxylase activity |
| **CIT** | Citrate utilization |
| **H<sub>2</sub>S** | Hydrogen sulfide production |
| **URE** | Urease activity |
| **TDA** | Tryptophan deaminase |
| **IND** | Indole production from tryptophan |
| **VP** | Acetoin production (Voges-Proskauer test) |
| **GEL** | Gelatinase activity |
| **GLU** | Glucose fermentation |
| **MAN** | Mannitol fermentation |
| **INO** | Inositol fermentation |
| **SOR** | Sorbitol fermentation |
| **RHA** | Rhamnose fermentation |
| **SAC** | Sucrose fermentation |
| **MEL** | Melibiose fermentation |
| **AMY** | Amygdalin fermentation |
| **ARA** | Arabinose fermentation |

#### Genetic characterisation

The genetic characterisation of the strains was done in the most promising bacteria to confirm their identification by biochemical characterization.

Genomic DNA was extracted from the cultures received using the Quick-DNA™ Fungal/Bacterial Miniprep Kit (Zymo Research, Catalogue No. D6005). The 16S target region was amplified using OneTaq® Quick-Load® 2X Master Mix (NEB, Catalogue No. M0486) with the primers presented in Table 1. The PCR products were run on a gel and cleaned up enzymatically using the EXOSAP method. The extracted fragments were sequenced in the forward and reverse direction (Nimagen, BrilliantDye™ Terminator Cycle Sequencing Kit V3.1, BRD3-100/1000) and purified (Zymo Research, ZR-96 DNA Sequencing Clean-up Kit™, Catalogue No. D4050). The purified fragments were analysed on the ABI 3500xl Genetic Analyzer (Applied Biosystems, ThermoFisher Scientific) for each reaction for every sample, as listed in Section 1. BioEdit Sequence Alignment Editor version 7.2.5 was used to analyse the. ab1 files generated by the ABI 3500XL Genetic Analyzer and results were obtained by a BLAST search (NCBI)[31].

**Table 1:**
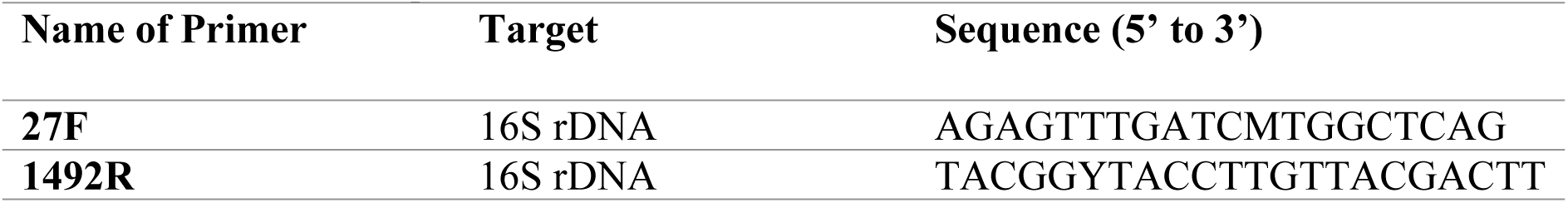
16S Primers sequences.

### Statistical analysis

All the raw data were recorded on an MS Excel 2010 sheet and Graph Pad prism 8.0. They were used to create tables, charts and calculations. the comparison between the assay was done using the fisher test at p<0.05.

The evolutionary history was inferred using the Neighbour-Joining method [32]. The optimal tree is shown. The tree is drawn to scale, with branch lengths in the same units as those of the evolutionary distances used to infer the phylogenetic tree. The evolutionary distances were computed using the Kimura 2-parameter method [33] and are in the units of the number of base substitutions per site. This analysis involved 34 nucleotide sequences. All ambiguous positions were removed for each sequence pair (pairwise deletion option). There were a total of 1804 positions in the final dataset. Evolutionary analyses were conducted in MEGA11 [34].

## Results and discussion

### Preparation of crude extracts from the endophytes isolated from *Cola nitida*

#### Isolation of endophytic bacteria

Endophytes were isolated from the leaves, trunk, branches and roots of *Cola nitida* using plant tissue culture on solid media and continuous culturing until pure colonies were observed. The figure 1 below shows that all *Cola nitida* organs were colonized by endophytic bacteria, with branches harboring the highest proportion (30.4%), followed by root bark (26.0%) and roots (20.0%). Trunk tissues showed moderate colonization (17.4%), while leaves had the lowest occurrence (6.5%). In total, 32 morphologically distinct bacterial isolates were recovered, indicating that both aerial and subterranean tissues serve as important niches for endophytic bacteria. These findings are consistent with previous reports that woody stems and root-associated tissues often support greater endophytic diversity due to their vascular structure and nutrient-rich microenvironments [35, 36]

**Figure 1.**
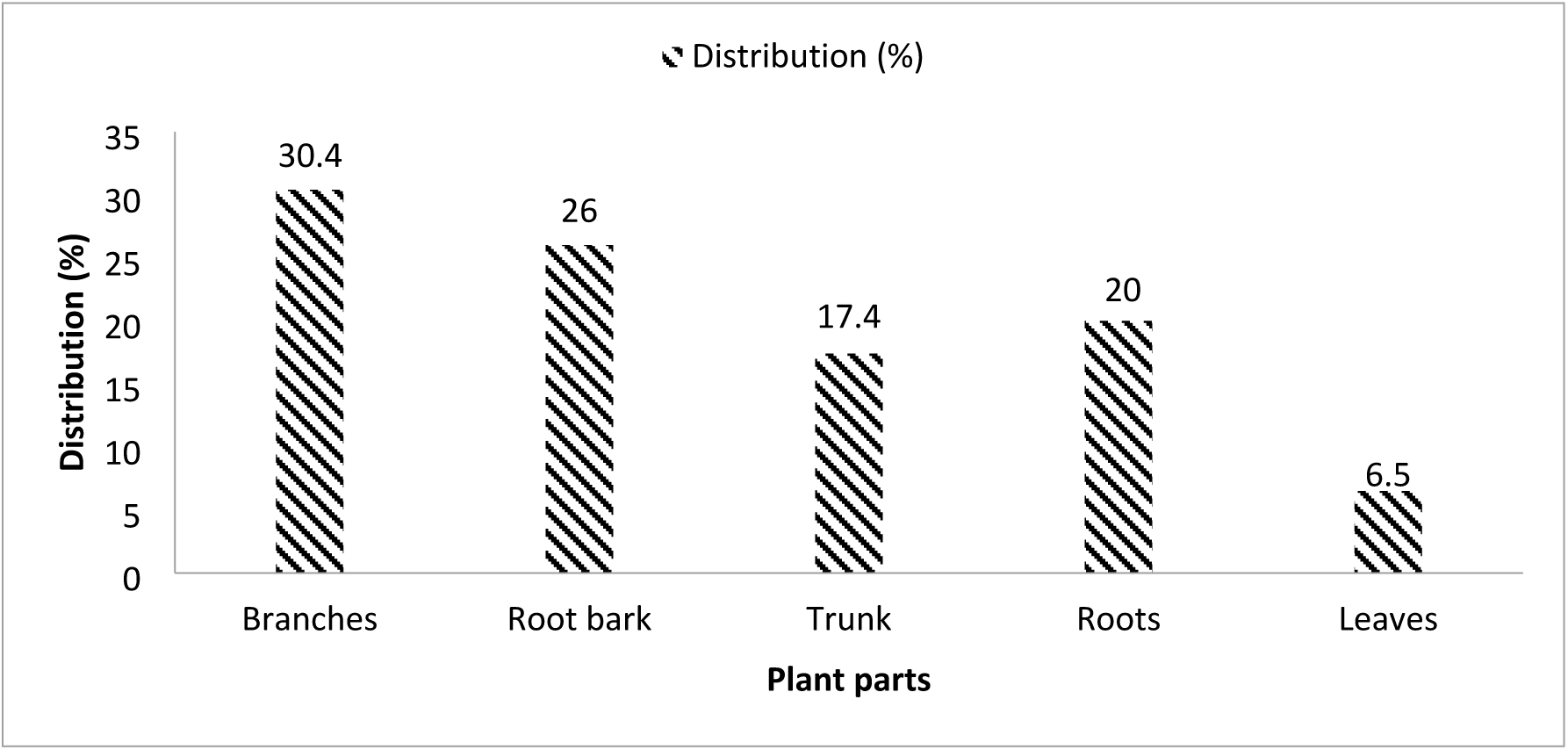
Distribution (%) of endophytic bacteria in different plant parts (*Cola nitida)*

### Assessment of Larvicidal Potency of endophytic bacteria

#### Preliminary screening of endophytic bacteria extract

After the larvicidal screening and based on the observed mortality, endophytes that showed no or only moderate larvicidal activity were excluded from further analysis. Of the 32 strains tested, only six exhibited promising larvicidal effects and were therefore selected for subsequent investigations. These endophytes include NPMRU 6211; NPMRU 6600; NPMRU 6164; NPMRU 6501; NPMRU 6186-1; NPMRU 6512-1

#### V.2.2 Assessment of the optimal dose of selected endophytic bacteria extracts

The data in Figure 2 shows that the more concentrated the extract, the deadlier it got for *Anopheles gambiae* larvae. At 0.5 MCF, kill rates stayed modest, somewhere between 10% and 40%, with isolates 6600 and 6501 topping out at 40%. Push the concentration up to 2 MCF, though, and things change fast, every extract jumps to 70-80% mortality. At 4 MCF, four isolates, 6211, 6600, 6501, and 6186-1, wiped out the larvae entirely. Therefore, 2 MCF looks like the sweet spot for real-world use, while 4 MCF shows just how lethal these extracts can get.

**Figure 2.**
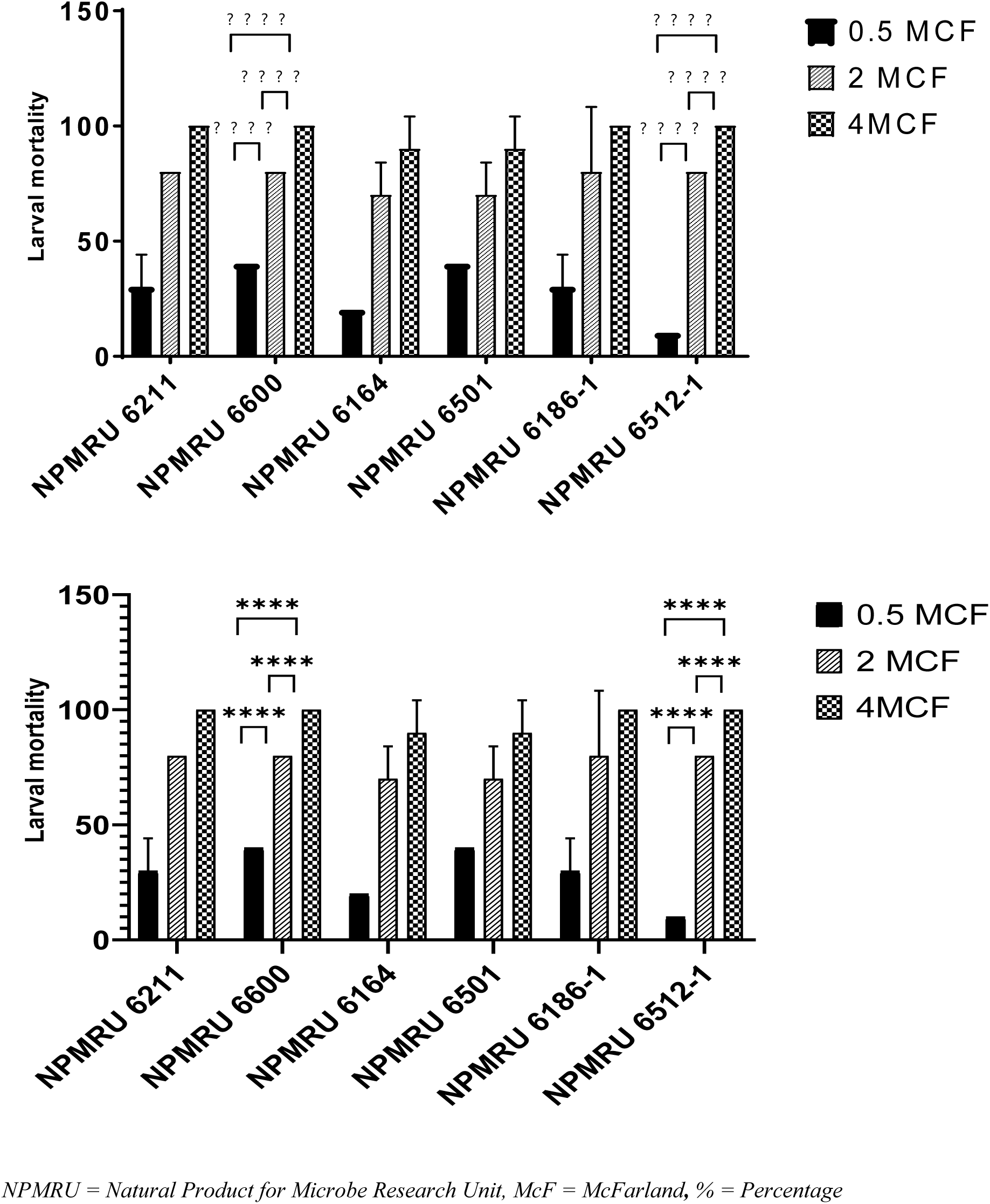
Larval mortality (%) of *Anopheles gambiae* exposed to six endophytic extracts at different McFarland scale

To the best of our reading, we did not find a report on the use of endophyte use in a biological fight against larva. However, this finding goes in the same line as that of Derua *et al.,* revealed that *Bacillus thuringiensis var. israelensis* and *Bacillus sphaericus* produce equally secondary metabolites that kill Anopheles gambiae on dose-dependent manner. Many other bacteria have shown the same trend.

What’s striking is where the active isolates came from, different organs of Cola nitida, suggesting potent larvicidal microbes hide throughout the plant, with roots and woody tissue often proving especially rich, and worth digging into further.

#### V.2.2 Assessment of the lethal inoculum size of selected endophytic bacteria

As illustrated in the figure 3 below, analysis revealed that NPMRU 6512-1 exhibited the highest larvicidal potency, with an LC₅₀ value of 0.747UFC/ml, indicating its ability to kill 50% of larvae at the inoculum size tested. This was followed by NPMRU 6164 and NPMRU 6186-1, both with LC₅₀ values of 1.384, showing strong larvicidal potential. NPMRU 6211 and NPMRU 6600 demonstrated moderate effectiveness, each with an LC₅₀ of 1.621. In contrast, NPMRU 6501 displayed the least potency, with an LC₅₀ of 2.116, suggesting a comparatively weaker effect. These results indicate that NPMRU 6512-1, 6164, and 6186-1 are the most promising candidates respectively for further investigation as larvicidal agents

**Figure 3.**
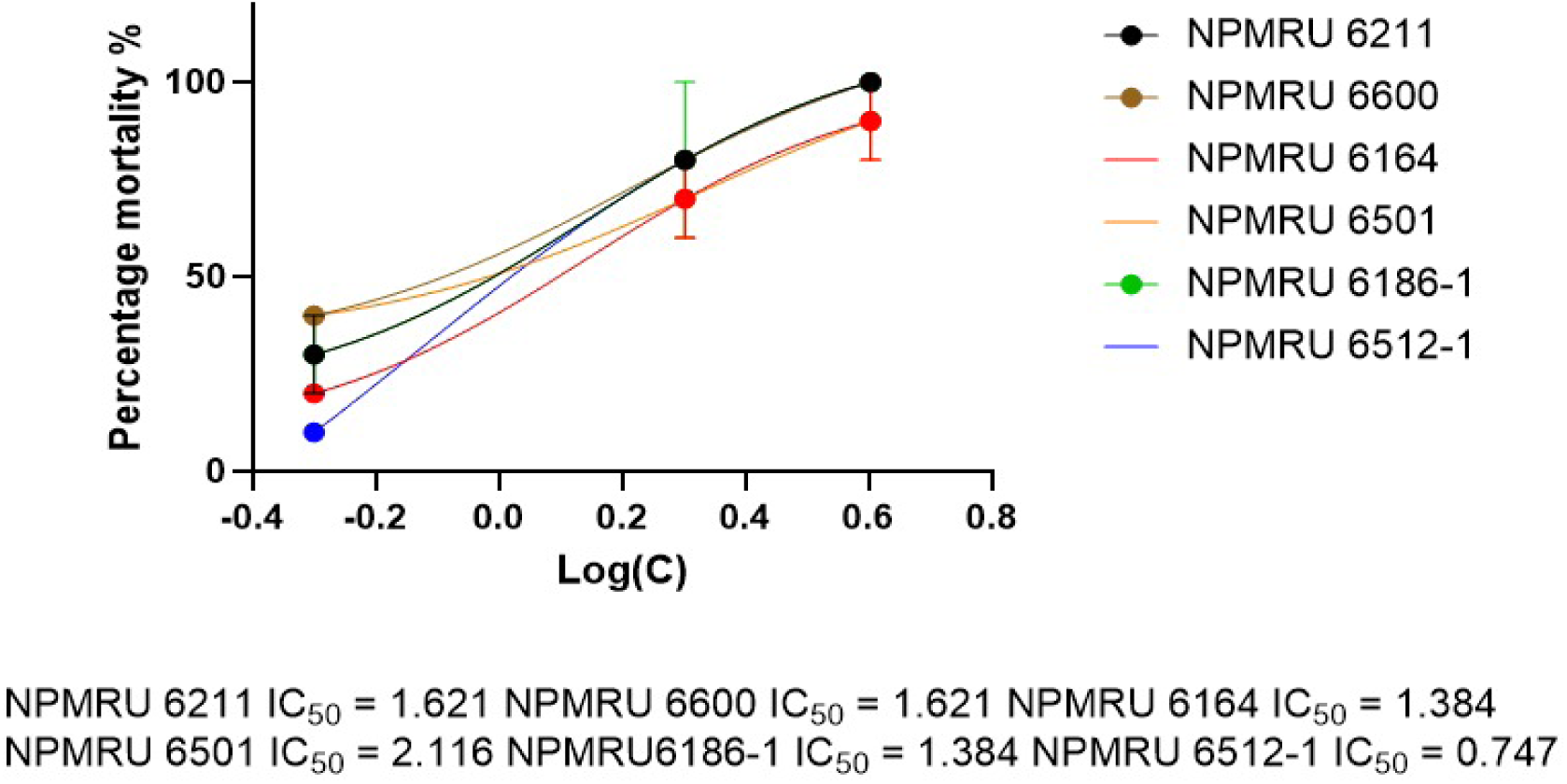
Larvicidal potency of bacterial extracts based on LC50

To the best of our knowledge, this is the first study using endophytic bacteria-larvae direct confrontation. However, the result obtained goes in the in the same line with what was reported by Gnambani *et al.,* [37] on *Chromobacterium anophelis sp.* In fact, these authors reported that the *C. anophelis* are killing *Anopheles coluzzii,* a member of the *An. gambiae* species complex both by robust colonization and pathogenic capacity or producing secondary metabolites. On the other hand many bacterial extract are producing compounds with larvicidal activity, employing mechanisms that range from highly specific receptor-mediated pore formation (Bti Cry toxins) and receptor-mediated apoptosis (*L. sphaericus* Bin toxin) to broad-spectrum midgut destruction (Csp_P)[17, 38], neurotoxic paralysis (PMP1), enzymatic degradation of structural barriers (chitinases from Pseudomonas and *B. laterosporus*), metabolic poisoning (HCN from *Chromobacterium*)[39], DNA damage and oxidative stress (sterigmatocystin from endophytic *Podospora*)[21].

### Environmental toxicity evaluation using the *Lemna minor* inhibition test

*Cola nitida* endophytic extracts were tested on *Lemna minor* at 2 McF and 4 McF, with observations on Days 0, 3, and 7 for frond development and appearance. At 2 McF all treated plants stayed green and healthy; each extract produced ≥3 fronds by Days 3 and 7, and controls also showed normal growth. At 4 McF phytotoxicity appeared (Table 2) : extract 6211 caused frond loss and severe chlorosis by Day 7; extracts 6600 and 6186-1 produced <2 fronds with tissue damage; extracts 6164 and 6512-1 destroyed plants (0 fronds). Results indicate dose-dependent toxicity at 4 McF.

**Table 2:** Aspect and number of *Lemna minor* fronds in 4MCF at day 0, day 3 and day 7.

| NPMRU | A0 | F0 | A3 | F3 | A7 | F7 |
| --- | --- | --- | --- | --- | --- | --- |
| **Control 1** | Green | 2 | Green | $\geq 3$ | Green | $\geq 3$ |
| **Control 2** | Green | 2 | Green | $\geq 3$ | Green | $\geq 3$ |
| **Control 3** | Green | 2 | Green | $\geq 3$ | Green | $\geq 3$ |
| **6211** | Green | 2 | Pale | 2 | Very chlorotic and partially destroyed | 2 |
| **6600** | Green | 2 | Pale | 2 | Very chlorotic and partially destroyed | 2 |
| **6164** | Green | 2 | Pale | 2 | Completely destroyed | 0 |
| **6501** | Green | 2 | Slight chlorosis | 2 | Chlorotic | 2 |
| **6186-1** | Green | 2 | Pale | 2 | Very chlorotic and partially destroyed | 2 |
| **6512-1** | Green | 2 | Pale | 2 | Completely destroyed | 0 |

Key: % = Percentage; NPMRU = Natural Product for Microbe Research Unit; A = Frond Aspect, F = Frond Number; ≥ = Greater or equals to

According to WHO, larvicides for field use must be both effective and low in ecological impact, achieving target mortality at doses that spare non-target organisms. *Cola nitida* endophytes were tested on *Lemna minor* at 2 MCF and 4 MCF over seven days, recording frond number and appearance on Days 0, 3, and 7. At 2 MCF, plants stayed green and healthy; frond counts remained ≥3 and matched controls, showing no visible phytotoxicity. By contrast, 4 MCF produced clear, dose-dependent damage. Some extracts (6211, 6600, 6186-1) caused chlorosis and reduced frond counts below two by Day 7; others (6164, 6512-1) completely destroyed plants, leaving no fronds. One extract (6501) induced only mild chlorosis.

These outcomes indicate a narrow safety window: 4 MCF is phytotoxic, whereas 2 MCF corresponds to a No Observed Effect Concentration in this assay. This pattern aligns with literature showing L. minor tolerates low stressors but displays frond loss, chlorosis, and structural damage at higher exposures. For field application, these data support selecting 2 MCF as the environmentally safer concentration for *Cola nitida* endophytic larvicides, balancing larval efficacy with protection of aquatic non-target plants.

### Characterisation and identification of most active endophytic bacteria

Some selected endophytic bacteria, NPMRU6600 (rootbark) and NPMRU6211 (trunk), were examined macroscopically. NPMRU6600 forms circular, well-defined colonies with a dark brown center fading toward the margins; the surface shows radial branching, a rough, dry texture, and lobulate to filamentous edges resembling hyphae. Elevation is flat to slightly raised and colonies are opaque.

**Figure 4.**
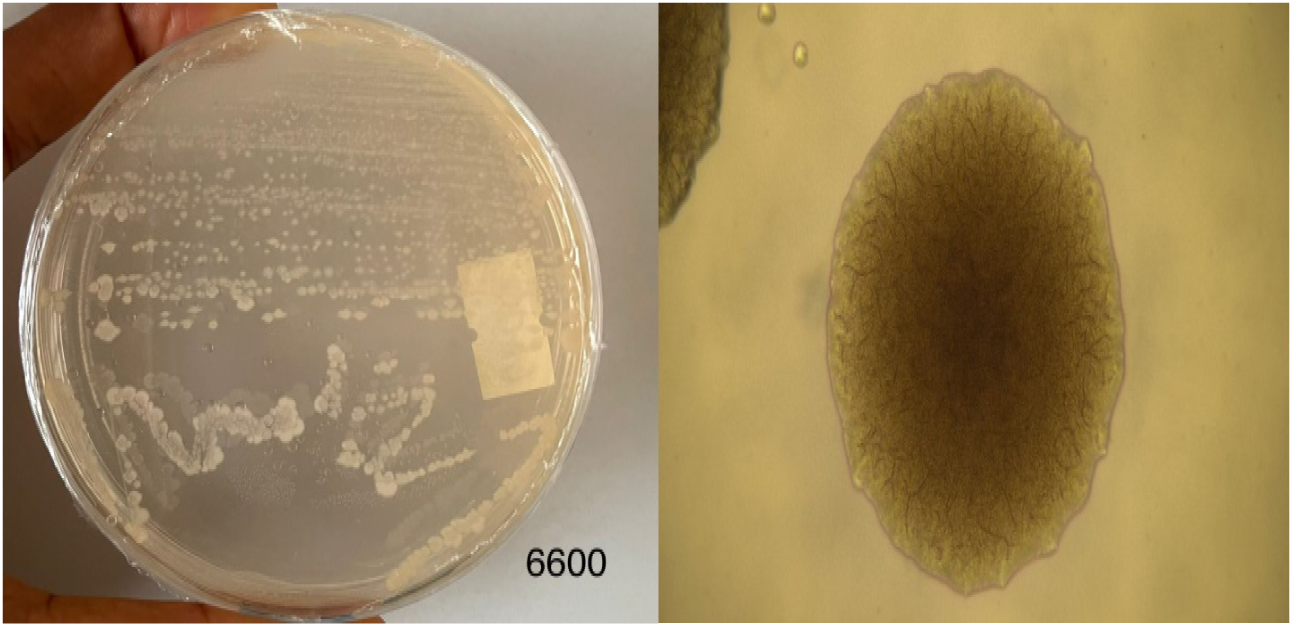
Macroscopic aspects of endophyte NPMRU6600

NPMRU6211 produces irregular, spreading colonies with lobate margins, pale beige to light brown color (darker centrally), and a smooth, moist surface. Elevation is flat to slightly raised, edges are diffuse (suggesting motility or surfactant activity), and texture appears soft or mucoid with translucent outer zones.

**Figure 5.**
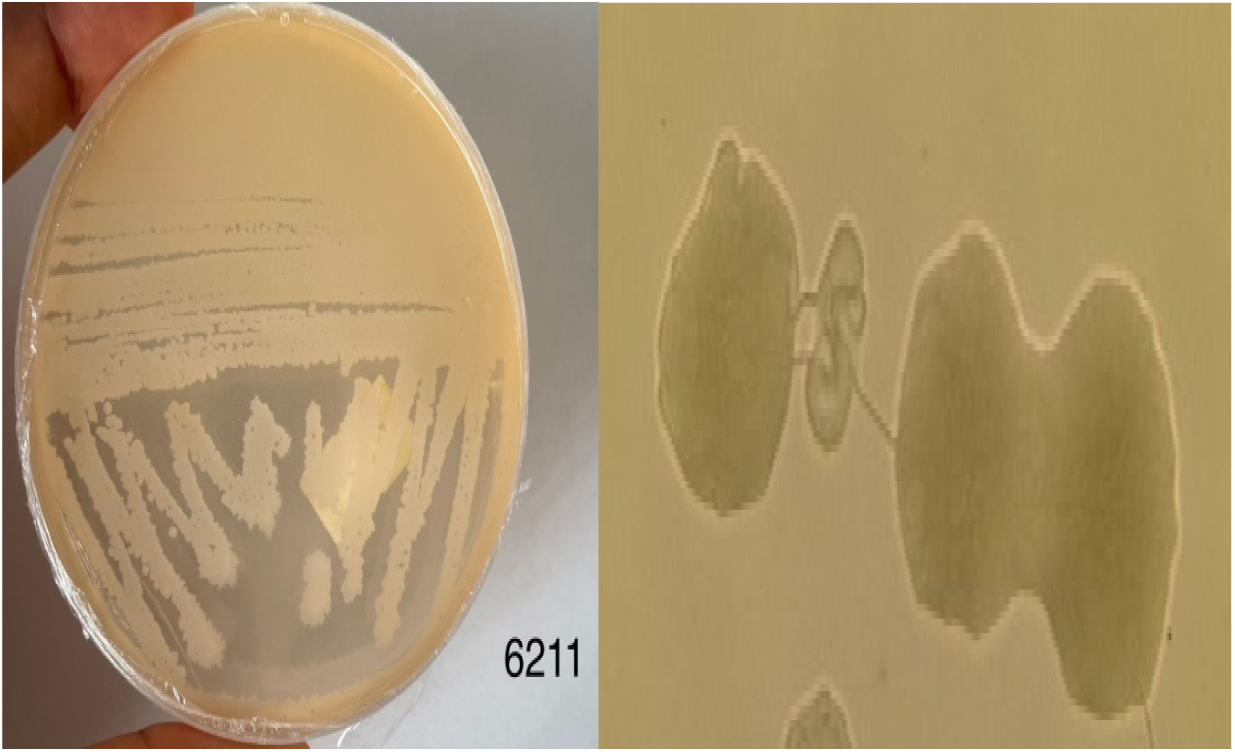
Macroscopic aspects of endophyte NPMRU6211

API 20E biochemical testing (interpreted via API gallery) yielded close matches to *Photobacterium damselae* (isolates 6164, 6211), *Salmonella enterica* spp. Arizonae (6600, 6501), and *Pseudomonas* spp. (6512-1, 6186-1), with 96–99.99% similarity supported by key markers. These data indicate consistent endophytic communities in Cola nitida.

**Table 3:**
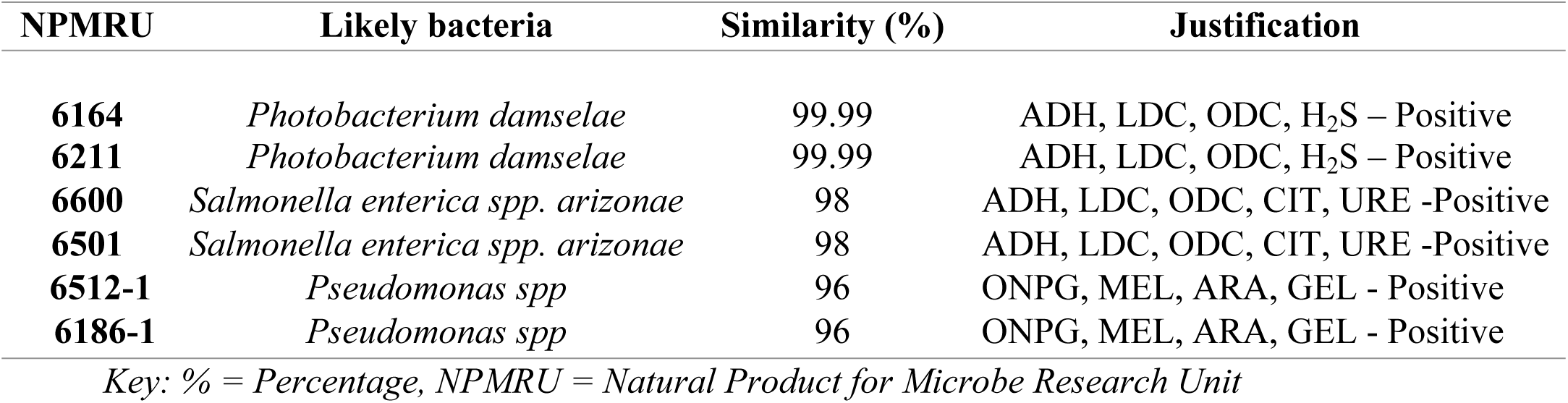
Endophytic bacteria identification using API 20E Biochemical gallery.

API 20E biochemical analysis identified three distinct bacterial genera within the endophytic collection (Table 3). *Photobacterium damselae* (NPMRU6164, NPMRU6211) demonstrated 99.99% sequence similarity, whereas *Salmonella enterica* subsp. *arizonae* (NPMRU6600, NPMRU6501) and *Pseudomonas* spp. (NPMRU6512-1, NPMRU6186-1) exhibited 98% and 96% identity respectively. Characteristic biochemical profiles substantiated all taxonomic assignments. Isolation of *Salmonella* and *Pseudomonas* as plant endophytes is consistent with documented occurrence of these genera in medicinal plants and their recognized antimicrobial and entomopathogenic potential[35, 36]. The marine-associated *Photobacterium* identification stands out and warrants deeper ecological investigation.

### Genetical identification

**Table 4:**
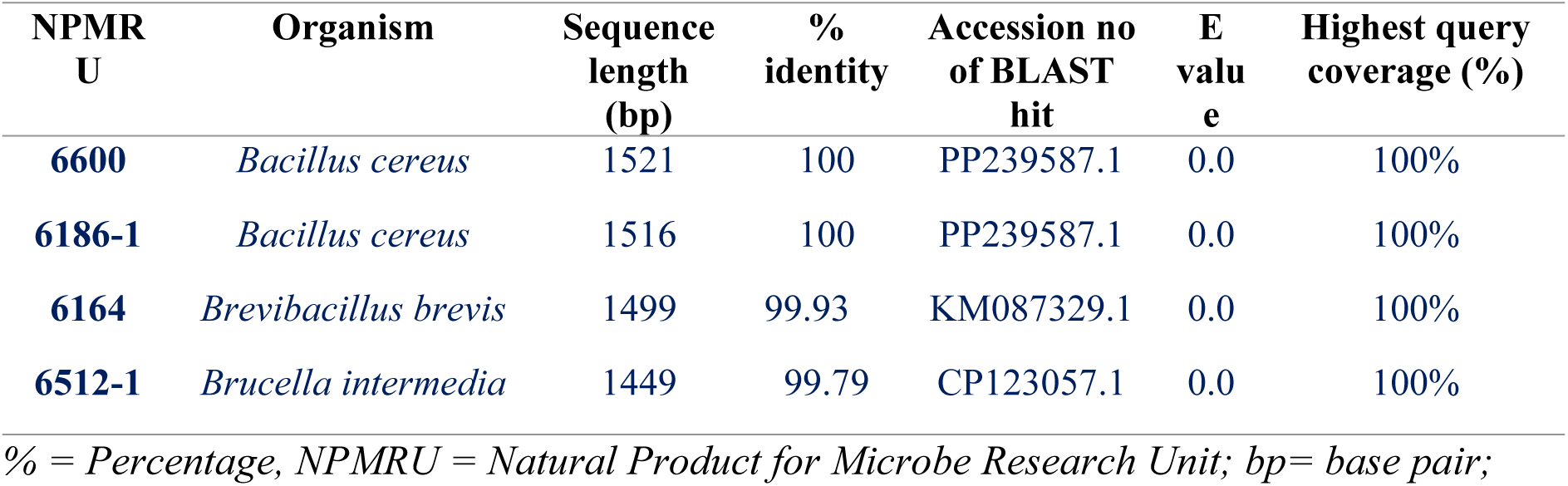
Endophytic bacteria identification using 16S rRNA gene sequencing and BLAST analysis.

16S rRNA gene sequencing and BLAST analysis successfully identified four distinct bacterial endophytes with high phylogenetic confidence (table 4). Two isolates (NPMRU6600, NPMRU6186-1) were assigned to *Bacillus cereus* with 100% sequence identity. NPMRU6164 clustered with *Brevibacillus brevis* (99.93%), while NPMRU6512-1 aligned to *Brucella intermedia* (99.79%). All sequences achieved maximum query coverage with E-values of 0.0, confirming robust taxonomic assignments. Recovery of *Bacillus* and *Brevibacillus* species from plant tissue aligns with their documented endophytic occurrence.

Across this isolate set, PCR-based 16S rRNA gene sequencing and API 20E biochemical profiling arrived at fundamentally different genus-level assignments not in marginal cases, but consistently and in opposite directions. The divergence is most sharply illustrated by NPMRU 6600, which API 20E assigned to *Salmonella enterica* subsp. *arizonae*at 98% similarity, yet resolved by PCR as *Bacillus cereus* with 100% sequence identity across a 1521 bp read, complete query coverage, and an E-value of 0.0 against accession PP239587.1. A match of that statistical quality is unambiguous. The biochemical signals that drove the API 20E (ADH, LDC, ODC, CIT, and URE positivity) almost certainly reflect convergent metabolic activity rather than true phylogenetic affinity, a phenomenon well documented in *Bacillus* species, which can express decarboxylase and urease activities under standard incubation conditions and thereby generate false-positive matches in galleries calibrated for Gram-negative enterics [40, 41].

The same pattern recurs with NPMRU 6186-1 and NPMRU 6512-1, both placed by API 20E within *Pseudomonas* spp. at 96% similarity. PCR data resolved them as *Bacillus cereus* and *Brucella intermedia*, respectively. These two bacteria are Firmicute and an Alphaproteobacterium with no meaningful phylogenetic overlap. Their co-assignment by API 20E underscores what Janda & Abbott[42] described as the intrinsic limitation of phenotypic gallery systems: pattern-matching against a biochemical database cannot substitute for sequence homology when isolates fall outside the taxon set the system was trained on. The *Photobacterium damselae* assignment for NPMRU 6164 at 99.99% biochemical similarity, subsequently overturned by PCR identification as *Brevibacillus brevis*, extends this argument further. These two organisms span opposing domains of bacterial phylogeny — one a Gram-negative marine Gammaproteobacterium, the other a Gram-positive endospore-former. Yet their ADH, LDC, ODC, and H₂S reaction profiles converged enough to confound a system never designed for plant-associated endophytes.

Taken together with the broader literature on culture-independent microbial identification [42, 43] these data reinforce the argument that endophyte collections from tropical medicinal plants routinely harbor organisms whose biochemical diversity exceeds the reference frame of enteric-focused phenotypic systems. PCR delivered sequence lengths of 1449–1521 bp, identity values of 99.79–100%, and uniform E-values of 0.0 an internal consistency that API 20E similarity scores of 96–99.99% cannot replicate, because those scores carry a categorically different epistemic weight.

## Conclusion

Bacterial endophytes show genuine promise as greener alternatives to synthetic larvicides, biodegradable, ecologically safer, yet biologically potent. With insecticide resistance spreading through *Anopheles* populations, these active isolates offer a credible starting point for next-generation biolarvicidal formulations.

## Akcnowledgements

The authors thank AREF for funding PCR-based identification of the bacterial endophytes via fellowship AREF-316/332-HZOU-F-CO1014 awarded to Dr Hzounda Fokou Jean Baptiste.

